# Comparison of personality structures between methodologies: personality ratings and behavioural observations in Japanese macaques (*Macaca fuscata*)

**DOI:** 10.1101/2022.09.30.510262

**Authors:** Kosho Katayama, Masayuki Nakamichi, Kazunori Yamada

**Author notes:** Corresponding author: (KK).

## Abstract

In this study, we aimed to clarify the structure of personality of Japanese macaques (*Macaca fuscata*) using two methods, personality ratings and behavioural observation, which are frequently used in primate personality studies. Principal component analysis of personality ratings sampled in two waves that were assessed with questionnaires (Stevenson-Hinde and Zunz questionnaire in Wave 1 and Hominoid Personality Questionnaire in Wave 2) revealed four components for each wave: Dominance, Aggressiveness, Dependency, and Component4 and Dominance, Aggressiveness, Sociability, and Activity. By calculating the correlation coefficients for principal component loadings of personality ratings, we compared the personality structure of other primates from previous studies with that of Japanese macaques. Based on these results, we propose that the factors common to macaques and capuchin monkeys fall into the following four categories of Dominance, which correlates with dominance rank; Excitability, which expresses aggression and neuroticism; Sociability, which indicates how actively they try to interact with others; and Activity, which expresses exploratory behaviour and openness. Analysis of data from behavioural observation extracted four components: Kin-biased-approaching, Grooming-diversity, Activity, and Aggressiveness. Principal component scores were calculated from each rating and behavioural observation data, and correlation coefficients were used to examine whether the same personality traits were found in both these methods. In this analysis, the raters did not collect behavioural data to ensure that data from both these methods did not overlap. No identical personality traits were found between the two methods. This result may indicate that there was no specific personality structure in Japanese macaques and that each method assessed different aspects of behavioural patterns pertaining to animal behaviour. It is difficult to assume from this study the cross-situational consistency conventionally assumed by personality psychology, and the results suggest that individuals vary in the way their behaviour changes in response to their environment, that is, coherence.

## Introduction

Consistent individual differences in behaviour between individuals of the same species are referred to as ‘personality’ and have recently received attention from biologists and psychologists [1,2]. One of the representative theories pertaining to personality is the trait theory, which states that personality structure is composed of several factors, called traits, and its underlying dimensions are usually identified through statistical methods such as factor analysis. For example, the Big Five model attempts to express human personality using five factors [3]. As traits have been interpreted as being related to internal mechanisms of individuals, temporal stability and cross-situational consistency are prerequisites for establishing personality traits of human and non-human primates. Several studies of non-human primates have demonstrated moderate temporal stability of personality [4] and consistency of behaviour under certain circumstances [5,6], whereas cross-situational consistency in personality traits through diverse situations has remained a controversial topic in human and non-human primate personality theory [7–9].

There have been numerous studies that have revealed the social behaviour of non-human primates. These include descriptions of social structure, such as demographic composition, kin relationships, and factors that shape individual association patterns within groups [10,11]. In Macaques and baboons species, grooming and proximity behaviours are more frequent among kin-biased individuals than among non-kin individuals [12–14]. Macaca species can be classified into two groups according to their dominance style: despotic and egalitarian [15,16]. Grooming is more kin-biased in despotic macaque species than in egalitarian species and is usually directed up high-ranking individuals [12,17]. Conversely, some individuals in a despotic macaque group groom non-kin or lower-ranking individuals more frequently than kin or higher-ranking ones [18]. Several studies have examined how individual differences in macaques are represented in the context of trait theory [19–21]. For example, [20]factor analysis performed on behavioural data of wild chacma baboons expressed its personality structure through three factors labelled Aloof, Loner, and Nice [21]. Nice was not related to dominance rank but to fitness indices such as composite sociality indices. Many studies have examined primate personality in captive populations. However, only a few have attempted to capture overall individual differences in social and non-social traits through behavioural observations of wild populations. Further research is needed to determine how social structures, such as dominance rank relationships and number of kin, shape temporal stability and cross-situational consistency in personality.

Previous studies have attempted to elucidate the latent personality structure of non-human primates using questionnaires and personality ratings (PRs) [22–24], but there have been few studies in this regard on Japanese macaques. The provisioned Japanese macaque group on Koshima Island, the wild population of Yakushima Island, and the zoo population in Italy [22] have been studied using a rating method, stating that the personality structure of these macaques can be expressed by four factors: dominance, openness, friendliness, and anxiety. The structure of personality composition between non-human primates was systematically compared using rating methods, and an evolutionary lineage of primate personality was proposed [25]. However, few studies have replicated this research [22,24,26,27], and several non-human primates, such as crested macaques (*Macaca nigra*), have not been included in them. Further research is required to validate evolutionary phylogeny of personality traits in non-human primates.

Previous studies have pointed out advantages of rating methods, such as capturing subtle individual differences and overall behavioural patterns [28]. The personality structure of rating data was associated with external criteria, such as immune response [29]. Traits extracted from personality ratings are also associated with behavioural data; for example, the trait ‘Sociable’ in rhesus macaques (*Macaca mulatta*) is associated with behavioural data such as lip-smack or groom [5]. There are also data that test construct validity in Chimpanzees (*Pan troglodytes*), showing that semantically similar traits and behavioural data are correlated, similar to the trait ‘dominance’ found by personality ratings being associated with submissive behaviour [30]. Applications of ratings is also expanding, with some studies attempting to use it for animal management [31]. However, other studies have pointed out that personality structure of ratings cannot reflect the component structure found from behavioural data [27,32]and that personality ratings have methodological issues [33–35], which refer to several kinds of biases derived from the raters’ abstract perception of behaviours and stereotypical beliefs about individuals [32,33].

Thus, the question of whether the same personality structure can be found in individuals of non-human primates, regardless of the methodology, is similar to the problem raised by the person-situation debate elucidated in Mischel’s argument [8]. According to the traditional trait theory, latent traits should lead to cross-situational consistency in behaviour and the same trait structure, regardless of methodology. Moreover, it was also argued that behaviour is strongly influenced by the situation and that the trait lacks cross-situational consistency. However, the question remains that how can these seemingly conflicting claims be reconciled. Some solutions to this person-situation debate have already been proposed [7,36–38], but it is not yet clear whether they can be applied to animal personality research. Recently, researchers have argued that personality traits are a result of expressing the interaction among certain sets of factors, such as individuals in quantification, occasions, spans of time, and situations [33,39,40]. Ethological research that records behaviour continuously through observation across a wide range of natural settings can provide useful data for testing the human trait theory. In this study, we aimed to elucidate the personality structure of wild Japanese macaques using both personality ratings and behavioural observation (BO) and compare the personality structures of Japanese macaques and other non-human primates, such as capuchin monkeys (*Sapajus apella*), for which no comparative study has been conducted so far. We also examined whether the same personality structure could be found from both, personality ratings and behavioural observations, of provisioned Japanese macaques in wild populations.

## Materials and Methods

### Ethics statement

All experiments were conducted in accordance with the Regulations on Animal Experimentation at Osaka University, Japan, and were approved by the Animal Research Committee of the Graduate School of Human Sciences, Osaka University (No. 24-2-0). As the study site, Iwatayama Monkey Park, Kyoto, Japan, is privately owned, observations were made with the permission of the management.

### Study site and subjects

The study subjects were 35 Japanese macaques (S1 Table) from a free-ranging, provisioned group (the Arashiyama group) living in Iwatayama Monkey Park (35°00′N, 135°67′E), Arashiyama, Kyoto, Japan. The study was conducted between July 2012 and October 2015. In August 2013, the group comprised 126 individuals, including 10 adult males (≥ 5 years old), 90 adult females (≥ 5 years old), 19 immatures (aged 1–4 years), and 7 infants (aged <1 year). Kin-relations through maternal lines and ages of individuals in the group were known since the introduction of provisioning in 1954 (see [41,42] for detailed information on this group). Dominance rank relationships between subjects were determined via observation of dyadic social interactions in the group. In the Arashiyama group, breeding season was from late September to March and the birth season, from April to August [43].

### Personality ratings and behavioural observation

Personality ratings were obtained by eight raters in two waves. The raters were park staff and researchers from a university, who had at least three months of work or study experience with this subject group. Five of the eight raters could identify all individuals in the group based on facial or behavioural characteristics. Wave 1 sample consisted of 30 adult females (mean age ± SD = 18.00 ± 3.44 years; S1 Table) rated between July and September 2012 by four raters. Fifteen subjects each were selected from among high and low-ranking individuals. Subjects were also selected from 67 females between 10 and 26 years of age, with maximum age variation. The questionnaire was adapted from Stevenson-Hinde and Zunz questionnaire [23] and consisted of 17 adjectives, in which 12 were corresponding descriptions, and five were additional adjectives (S2 Table contains the list of adjectives and their descriptions).

In Wave 2 sample, five males were rated in addition to the 30 females rated in Wave 1 between October and November 2015. During Wave 2, three out of the 30 females rated in Wave 1 had died, bringing the total to 27 females (mean age ± SD = 20.93 ± 3.58 years; S1 Table) and five males (mean age ± SD = 23.80 ± 6.14 years; S1 Table). In Wave 2, personality of the subjects was assessed using the Hominoid Personality Questionnaire (HPQ) consisting of 54 adjectives [22,25], which were scored on a seven-point Likert-type scale. In addition to these 54 items, an item used in Wave 1 was added to make a total of 55 items (S3 Table). There were five raters in Wave 2, but one rater did not complete the questionnaire, and therefore, their data were excluded from the analysis, leaving a final total of four raters including the first author. All raters were explicitly cautioned to not discuss their ratings with each other. All the questionnaires were translated from English to Japanese.

Behavioural observation was conducted by the first author (K.K.) between November 2013 and October 2015. Behavioural data from 32 individuals rated in Wave 2 personality ratings were collected using the focal animal sampling method [44]. Focal observations (20 min) were conducted from 0900 to 1730 h, when most macaques stayed near the provisioning site. Fifty-eight social and non-social behaviours that occurred in the daily routine of the Arashiyama group were recorded to cover a broad spectrum of behaviour using an all-occurrence sampling method or instantaneous sampling method with 30 s intervals (S4 Table). A wristwatch with an alarm (CASIO DW5600-1) and sampling sheets were used to record behaviour. Macaques were usually fed by the staff three times during the park’s open hours. No observations were made in this time, because the macaque density around the provisioning ground increased during the feeding periods. The subjects were observed in a predetermined random order. Before collecting data, all group members were identified based on their facial or behavioural characteristics. Maternal kin relations were defined as individuals with a k degree of relatedness ≥ 0.25 or more. That is, the mother-child, sister, and grandmother-grandchild relationships were defined as kin. The total observation time was 233 h (mean ± SD = 7.27 ± 0.61 h). All analyses were performed using R software (version 2.15.2 or 3.2.2) [45,46].

### Principal component analysis of personality ratings

Inter-rater reliability of each personality trait was calculated from the scores for 30 macaques in Wave 1 and 32 macaques in Wave 2. We used two intra-class correlation coefficients [47]: *ICC* (3, 1) and *ICC* (3, *k*). In Wave 1, *ICCs* were calculated from scores given by three raters. In Wave 2, two sets of *ICCs* were calculated given by four and three raters, respectively.

All item scores were standardised to a mean of zero with standard deviation being one. We then conducted a principal component analysis using principal procedure [48] and determined the number of components by examining the scree plot and conducting a parallel analysis [49]. We used both the varimax and promax procedures. The individual adjective scores were based on the mean scores calculated across ratings by the raters. We labelled each component by adjectives that had salient loadings (defined as 0.40) and other macaque personality literature [4,5,22,25]. To investigate the difference between Japanese macaque personality structure and that of other non-human primates, we examined Pearson’s correlation coefficient between component loadings based on Wave 2 scores by four raters and factor loadings based on other non-human primate personality structures rated by HPQ. For this analysis, we used results from previous studies on Barbary macaques (*Macaca sylvanus*), Assamese macaques (*Macaca assamensis*), Tonkean macaques (*Macaca tonkeana*), crested macaques [22], rhesus macaques [25], chimpanzees [50], orangutans (*Pongo sp*.; [51]), and capuchin monkeys [24]. To examine temporal stability of traits, Pearson’s correlation coefficient between the subject’s principal component score by three raters in Wave 1 and the scores by three raters in Wave 2 was calculated (N = 27). All six raters were different individuals to ensure that data from wave 1 and 2 did not overlap.

### Principal component analysis of behavioural measures

Principal component analysis was also performed on behavioural data. During the breeding season, the behaviour of males and oestrous females differs significantly compared to those during the non-oestrous season. Because the period during which the behavioural data were collected included the breeding season, the total number of observation targets were 32, but analysis was performed only on data from 27 females, excluding data of the five males. Behavioural data of females in oestrus were also excluded from this analysis. Interactions with individuals under three years of age were excluded from the data. Of the total 58 behavioural variables, 32 variables that were infrequent, strongly correlated, or observed only in a few specific individuals were excluded from analysis. Finally, 26 behavioural variables were selected to capture personality of the individual as widely as possible (S4 Table).

We also investigated whether the 26 behavioural measures were stable over time using *ICCs*. We divided the behavioural data collection period into two blocks and calculated frequencies of behaviours and indices describing the diversity of individuals in terms of social interaction partners in each of these time blocks separately for each subject (S1 Table). Period 1 was 67 days, from 20 November 2013 to 27 December 2014, and period 2 was 52 days, from 5 January 2015 to 30 October 2015. The total observation time was 80 h (mean per subject ± SD = 2.50 ± 0.38 h) in period 1 and 132 h (mean per subject ± SD = 4.12 ± 0.42 h) in period 2. Some variables were then transformed by the square root, square, or logarithm to achieve normal distribution and tested for *ICCs*. We considered only variables with statistical significance levels set at *p* < 0.05, that is, variables that showed temporal stability, were subsequently subjected to principal component analysis.

### Comparison of personality structure between personality ratings and behavioural observation

For each individual, principal component scores were calculated from the analysis of Wave 1 and Wave 2 ratings by three raters excluding the first author (K.K.), Wave 2 ratings by four raters including the first author (K.K.), and behavioural measuress, respectively. We examined relationships between these scores for 27 subjects and individual attributes of dominance rank, age, and number of kin using Pearson’s correlation coefficient. We examined whether the same personality structure of Japanese macaques was represented by personality ratings and behavioural observation. We calculated Pearson’s correlation coefficient between the principal component scores for 27 subjects from the rating data in Wave 2 and the behavioural data. In this analysis, the raters did not collect behavioural data.

## Results

### Personality structure of ratings in Wave 1 and Wave 2

We excluded four adjectives with *ICCs* less than zero (S5 Table) as unreliable measures [25] in Wave 1 ratings. In Wave 2, the principal component analysis required that the number of adjectives be less than the number of subjects, i.e. 32. Therefore, the top 31 reliable adjectives (S6 Table) were subjected to principal component analysis. Principal component analysis of the 13 reliable items in Wave 1 indicated four components with eigenvalues exceeding 1.00. The scree plot and parallel analysis suggested two components. As four factors have been extracted in previous studies on Japanese macaques [22], four components were also extracted in this study. Correlations between the varimax and promax-rotated principal component scores of the four components were high (*r* > 0.97); therefore, we decided to interpret personality structure and base the subject’s component scores on the varimax solution. We labelled the four Wave 1 components: Dominance_PR-1_, Aggressiveness_PR-1_, Dependency_PR-1_, and Component4_PR-1_ (Table 1). Because we found it difficult to interpret the fourth component and give it a specific name, we labelled it Component4_PR-1_. These four components accounted for 82 % of the total variance. The average absolute value of Pearson’s correlation coefficient between Wave 1 components and subject’s attributes was *r* = 0.34 (SD = 0.27, range 0.13–0.73) for subject’s dominance rank, *r* = 0.21 (SD = 0.08, range 0.12–0.30) for age, and *r* = 0.13 (SD = 0.08, range 0.03–0.23) for the total number of kin (Table 2).

**Table 1.** Factor loadings, eigenvalues, and cumulative contribution ratio of the four components calculated by principal component analysis (varimax rotation) in Wave 1 ratings (N = 30).

| Items | Components<sup>(1)</sup> |  |  |  |
| --- | --- | --- | --- | --- |
|  | Dom<sub>PR-1</sub> | Agg<sub>PR-1</sub> | Dep<sub>PR-1</sub> | Co4<sub>PR-1</sub> |
| Confidence | **0.90** | 0.33 | -0.14 | 0.05 |
| Fearful | **-0.84** | -0.10 | 0.04 | 0.08 |
| Solitary | **-0.87** | -0.22 | 0.09 | 0.12 |
| Sociable | **0.79** | 0.32 | 0.31 | 0.12 |
| Subordinate | **-0.66** | 0.31 | **0.43** | -0.15 |
| Fond of the opposite sex | **0.69** | 0.16 | **0.41** | -0.18 |
| Tolerant | **-0.43** | **-0.81** | 0.03 | 0.00 |
| Aggressive | 0.24 | **0.93** | 0.02 | 0.13 |
| Excitable | 0.11 | **0.94** | 0.09 | 0.20 |
| Equable | -0.04 | **-0.89** | -0.10 | -0.20 |
| Dependent | 0.01 | 0.02 | **0.95** | 0.06 |
| Watch for a chance | 0.10 | 0.21 | -0.14 | **0.84** |
| Fond of infants | 0.29 | -0.22 | -0.30 | **-0.67** |
| Eigenvalues | 4.17 | 3.67 | 1.50 | 1.34 |
| Cumulative contribution | 0.32 | 0.60 | 0.72 | 0.82 |

Salient loadings $\geq |.4|$ are shown in **bold**.
<sup>(1)</sup> Dom<sub>PR-1</sub> = Dominance<sub>PR-1</sub>, Agg<sub>PR-1</sub> = Aggressiveness<sub>PR-1</sub>, Dep<sub>PR-1</sub> = Dependency<sub>PR-1</sub>, and Co4<sub>PR-1</sub> = Component4<sub>PR-1</sub>

**Table 2.** Pearson correlation coefficients (*r*) and *p*-values between components and individual dominance rank, age, and the number of kin.

|  | Rank |  | Age |  | Number of kin |  |
| --- | --- | --- | --- | --- | --- | --- |
| Components | $r$ | $p$-value | $r$ | $p$-value | $r$ | $p$-value |
| ***Components in Wave 1 ratings (N = 30)*** |  |  |  |  |  |  |
| Dominance<sub>PR-1</sub> | -0.73 | $p < 0.001$ | 0.25 | 0.179 | 0.23 | 0.229 |
| Aggressiveness<sub>PR-1</sub> | -0.17 | 0.380 | 0.17 | 0.382 | 0.03 | 0.889 |
| Dependency<sub>PR-1</sub> | 0.32 | 0.085 | 0.30 | 0.101 | -0.13 | 0.481 |
| Component4<sub>PR-1</sub> | -0.13 | 0.490 | 0.12 | 0.543 | -0.14 | 0.445 |
| ***Components in Wave 2 ratings (N = 32)*** |  |  |  |  |  |  |
| ***4 raters*** |  |  |  |  |  |  |
| Dominance<sub>PR-2</sub> | -0.88 | $p < 0.001$ | 0.22 | 0.223 | -0.14 | 0.431 |
| Aggressiveness<sub>PR-2</sub> | -0.07 | 0.713 | -0.01 | 0.972 | -0.07 | 0.712 |
| Sociability<sub>PR-2</sub> | -0.09 | 0.606 | 0.01 | 0.974 | 0.51 | 0.003 |
| Activity<sub>PR-2</sub> | -0.14 | 0.449 | 0.85 | $p < 0.001$ | -0.14 | 0.436 |
| **3 raters** |  |  |  |  |  |  |
| Dominance<sub>PR-2-1</sub> | 0.87 | $p < 0.001$ | 0.34 | 0.056 | -0.06 | 0.755 |
| Aggressiveness<sub>PR-2-1</sub> | -0.16 | 0.386 | -0.22 | 0.223 | -0.16 | 0.391 |
| Sociability<sub>PR-2-1</sub> | -0.01 | 0.964 | -0.08 | 0.676 | 0.46 | 0.008 |
| Activity<sub>PR-2-1</sub> | -0.11 | 0.552 | 0.60 | $p < 0.001$ | -0.26 | 0.158 |
| **Components of behavioural measure (N = 27)** |  |  |  |  |  |  |
| Kin-biased-approaching<sub>BO</sub> | 0.17 | 0.407 | -0.22 | 0.260 | 0.74 | $p < 0.001$ |
| Grooming-diversity<sub>BO</sub> | -0.07 | 0.726 | -0.34 | 0.085 | -0.02 | 0.903 |
| Activity<sub>BO</sub> | 0.55 | 0.003 | -0.14 | 0.488 | -0.22 | 0.271 |
| Aggressiveness<sub>BO</sub> | 0.68 | $p < 0.001$ | -0.16 | 0.412 | -0.24 | 0.218 |

We conducted a principal component analysis of the 31 reliable items for the 32 subjects rated by four raters in Wave 2. The eigenvalue exceeded 1.00 for six components. The scree plot indicated there were four or six components, and parallel analysis suggested three components. In line with the Wave 1 analysis, we also extracted four components from Wave 2. Correlations between the four components’ varimax and promax-rotated principal component scores were high (*r* > 0.98). We decided to interpret the personality structure and base the subject’s component scores on varimax solution in the following analysis. We labelled the four components from Wave 2: Dominance_PR-2_, Aggressiveness_PR-2_, Sociability_PR-2_, and Activity_PR-2_ (Table 3). These four components account for 73 % of the total variance.

**Table 3.**
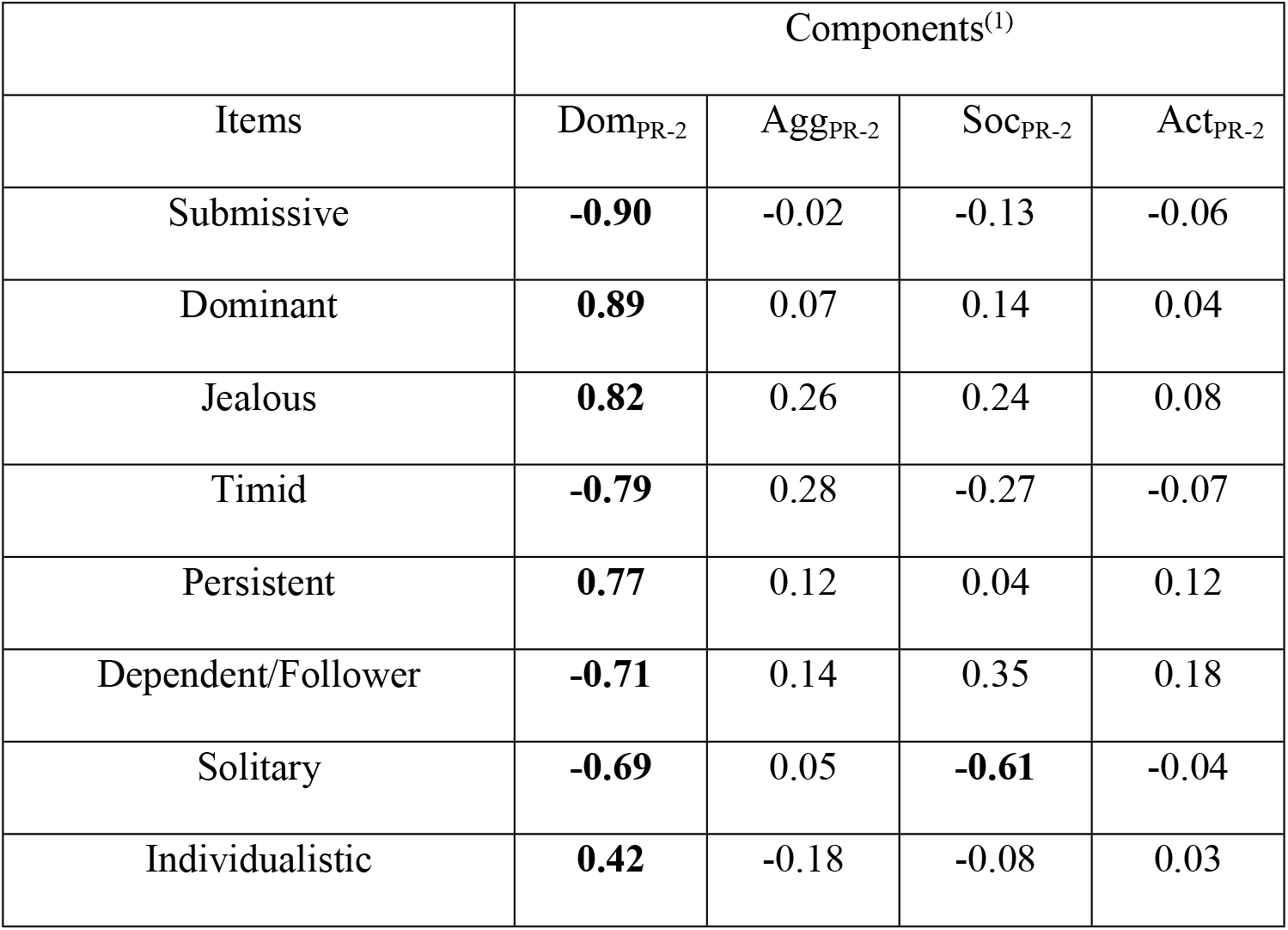
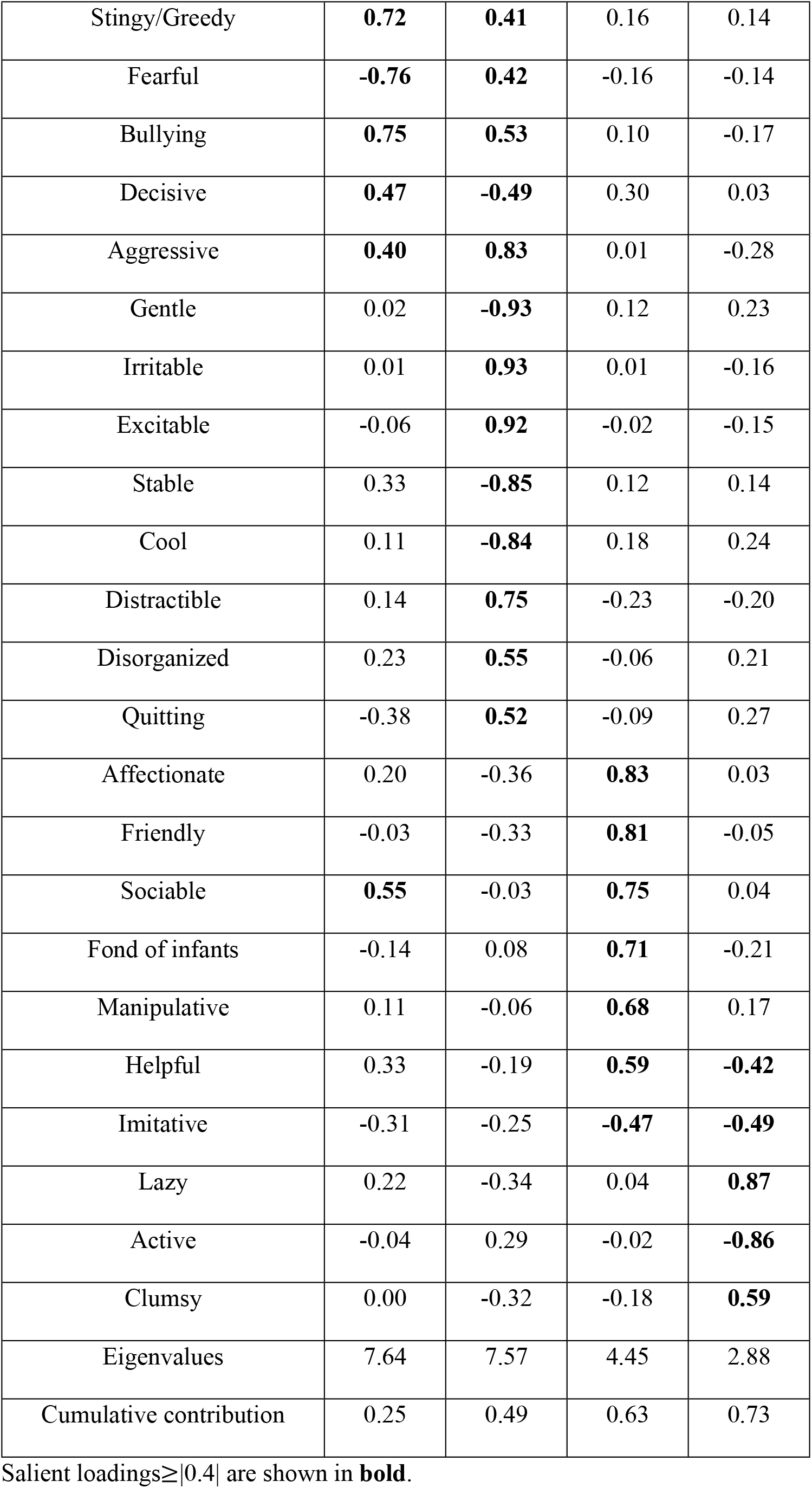
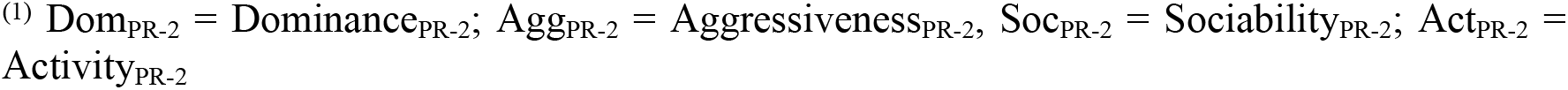
Factor loadings, eigenvalues, and cumulative contribution ratio of the four components calculated by principal component analysis (varimax rotation) in Wave 2 ratings (N = 32).

|  | Components<sup>(1)</sup> |  |  |  |
| --- | --- | --- | --- | --- |
| Items | Dom<sub>PR-2</sub> | Agg<sub>PR-2</sub> | Soc<sub>PR-2</sub> | Act<sub>PR-2</sub> |
| Submissive | **-0.90** | -0.02 | -0.13 | -0.06 |
| Dominant | **0.89** | 0.07 | 0.14 | 0.04 |
| Jealous | **0.82** | 0.26 | 0.24 | 0.08 |
| Timid | **-0.79** | 0.28 | -0.27 | -0.07 |
| Persistent | **0.77** | 0.12 | 0.04 | 0.12 |
| Dependent/Follower | **-0.71** | 0.14 | 0.35 | 0.18 |
| Solitary | **-0.69** | 0.05 | **-0.61** | -0.04 |
| Individualistic | **0.42** | -0.18 | -0.08 | 0.03 |
| Stingy/Greedy | **0.72** | **0.41** | 0.16 | 0.14 |
| Fearful | **-0.76** | **0.42** | -0.16 | -0.14 |
| Bullying | **0.75** | **0.53** | 0.10 | -0.17 |
| Decisive | **0.47** | **-0.49** | 0.30 | 0.03 |
| Aggressive | **0.40** | **0.83** | 0.01 | -0.28 |
| Gentle | 0.02 | **-0.93** | 0.12 | 0.23 |
| Irritable | 0.01 | **0.93** | 0.01 | -0.16 |
| Excitable | -0.06 | **0.92** | -0.02 | -0.15 |
| Stable | 0.33 | **-0.85** | 0.12 | 0.14 |
| Cool | 0.11 | **-0.84** | 0.18 | 0.24 |
| Distractible | 0.14 | **0.75** | -0.23 | -0.20 |
| Disorganized | 0.23 | **0.55** | -0.06 | 0.21 |
| Quitting | -0.38 | **0.52** | -0.09 | 0.27 |
| Affectionate | 0.20 | -0.36 | **0.83** | 0.03 |
| Friendly | -0.03 | -0.33 | **0.81** | -0.05 |
| Sociable | **0.55** | -0.03 | **0.75** | 0.04 |
| Fond of infants | -0.14 | 0.08 | **0.71** | -0.21 |
| Manipulative | 0.11 | -0.06 | **0.68** | 0.17 |
| Helpful | 0.33 | -0.19 | **0.59** | **-0.42** |
| Imitative | -0.31 | -0.25 | **-0.47** | **-0.49** |
| Lazy | 0.22 | -0.34 | 0.04 | **0.87** |
| Active | -0.04 | 0.29 | -0.02 | **-0.86** |
| Clumsy | 0.00 | -0.32 | -0.18 | **0.59** |
| Eigenvalues | 7.64 | 7.57 | 4.45 | 2.88 |
| Cumulative contribution | 0.25 | 0.49 | 0.63 | 0.73 |

Salient loadings $\geq |0.4|$ are shown in **bold**.
<sup>(1)</sup> Dom<sub>PR-2</sub> = Dominance<sub>PR-2</sub>; Agg<sub>PR-2</sub> = Aggressiveness<sub>PR-2</sub>; Soc<sub>PR-2</sub> = Sociability<sub>PR-2</sub>; Act<sub>PR-2</sub> = Activity<sub>PR-2</sub>

In Wave 2, principal component analysis was also performed on the adjectives by three raters other than the author, who collected the behavioural data. Similar to the analysis performed for the ratings by four raters, we performed principal component analysis on 31 adjectives with high inter-rater reliability and extracted four components. Because the correlation coefficient between the principal component scores by the four and three raters in Wave 2 was high (*r* > 0.73), we decided that the same components were obtained for both analyses and labelled them as Dominance_PR-2-1_, Aggressiveness_PR-2-1_, Sociability_PR-2-1_, and Activity_PR-2-1_ (S7 Table). These four components accounted for 75 % of the total variance.

In Wave 2 ratings by four raters, the average absolute value of Pearson’s correlation coefficient between the principal component score and subject’s attributes was *r* = 0.30 (SD = 0.39, range 0.07–0.88) for subject’s dominance rank, *r* = 0.27 (SD = 0.40, range 0.01–0.85) for age, and *r* = 0.22 (SD = 0.20, range 0.07–0.51) for the number of kin (Table 2). In Wave 2 ratings by three raters, the average Pearson’s correlation coefficient between the component score and subject’s attributes was *r* = 0.29 (SD = 0.39, range 0.01–0.87) for subject’s dominance rank, *r* = 0.31 (SD = 0.22, range 0.08–0.60) for age, and *r* = 0.24 (SD = 0.17, range 0.06–0.46) for the number of kin (Table 2). The traits with temporal stability from Wave 1 to Wave 2 (Table 4) were Dominance_PR-1_ (*r* = 0.66, *p* < 0.001) and Aggressiveness_PR-1_ (*r* = 0.57, *p* < 0.01).

**Table 4.** Pearson correlation coefficients (*r*) and *p*-values (N = 27) between the components of ratings from Wave 1 and Wave 2 (three-year temporal stability).

| Component | *Wave 2 ratings*<sup>(2)</sup> |  |  |  |  |  |  |  |
| --- | --- | --- | --- | --- | --- | --- | --- | --- |
|  | Dom<sub>PR-2</sub> |  | Agg<sub>PR-2</sub> |  | Soc<sub>PR-2</sub> |  | Act<sub>PR-2</sub> |  |
| | $r$ | $p$-value | $r$ | $p$-value | $r$ | $p$-value | $r$ | $p$-value |
| *Wave 1 ratings*<sup>(1)</sup> |  |  |  |  |  |  |  |  |
| Dom<sub>PR-1</sub> | 0.66 | $p < 0.001$ | 0.09 | 0.666 | 0.51 | 0.007 | 0.13 | 0.534 |
| Agg<sub>PR-1</sub> | 0.17 | 0.391 | 0.57 | 0.002 | 0.05 | 0.797 | 0.24 | 0.219 |
| Dep<sub>PR-1</sub> | -0.38 | 0.049 | 0.10 | 0.617 | 0.12 | 0.548 | 0.53 | 0.005 |
| Co4<sub>PR-1</sub> | 0.05 | 0.802 | 0.06 | 0.748 | -0.10 | 0.613 | -0.12 | 0.554 |

<sup>(1)</sup> Dom<sub>PR-1</sub> = Dominance<sub>PR-1</sub>, Agg<sub>PR-1</sub>, Aggressiveness<sub>PR-1</sub>, Dep<sub>PR-1</sub>, Dependency<sub>PR-1</sub>, and Co4<sub>PR-1</sub> = Component4<sub>PR-1</sub>
<sup>(2)</sup> Dom<sub>PR-2</sub> = Dominance<sub>PR-2</sub>, Agg<sub>PR-2</sub> = Aggressiveness<sub>PR-2</sub>, Soc<sub>PR-2</sub> = Sociability<sub>PR-2</sub>, and Act<sub>PR-2</sub> = Activity<sub>PR-2</sub>

### Comparison of personality structure between Japanese macaques and the other non-human primates

Table 5 shows Pearson’s correlation coefficient between principal component loadings of reliable adjectives in Wave 2 and the factor loadings of the other non-human primate studies. Dominance_PR-2_ was similar to factors Dominance or Confidence in other non-human primates; for example, Confidence of Barbary macaques (*r* = 0.84, *p* < 0.001). Aggressiveness_PR-2_ correlated with one to three factors in other primates (Table 5); for example, Aggressiveness of crested macaques (*r* = 0.83, *p* < 0.001) and Conscientiousness, Agreeableness, and Neuroticism of chimpanzees (*r* = −0.80, *p* < 0.001; *r* = −0.70, *p* < 0.001; *r* = 0.59, *p* < 0.01, respectively). Sociability_PR-2_ and Activity_PR-2_ also correlated with one to three factors in other non-human primates (Table 5); for example, Sociability_PR-2_ correlated with Friendliness of crested macaques (*r* = 0.57, *p* < 0.01) and Extraversion and Agreeableness of orangutans (*r* = 0.39, *p* < 0.05; *r* = 0.74, *p* < 0.001, respectively). Activity_PR-2_ was related to components such as Openness, Neuroticism, and Attentiveness of capuchin monkeys (*r* = −0.59, *p* < 0.001; *r* = −0.39, *p* < 0.05; *r* = −0.38, *p* < 0.05, respectively).

**Table 5.** Pearson correlation coefficients (*r*) and *p*-values between the principal component loadings of Wave 2 ratings and the factor loadings in other macaque species.

| Traits | *Japanese macaque (this study)* <sup>(1)</sup> |
| --- | --- |

|  | Dom<sub>PR-2</sub> |  | Agg<sub>PR-2</sub> |  | Soc<sub>PR-2</sub> |  | Act<sub>PR-2</sub> |  |
| --- | --- | --- | --- | --- | --- | --- | --- | --- |
|  | *r* | *p*-value | *r* | *p*-value | *r* | *p*-value | *r* | *p*-value |
| ***Japanese macaque (N = 30: [22])*<sup>(2)</sup>** |  |  |  |  |  |  |  |  |
| Dominance | 0.84 | *p* < 0.001 | 0.03 | 0.895 | 0.33 | 0.076 | -0.11 | 0.551 |
| Openness | 0.16 | 0.390 | 0.29 | 0.118 | 0.19 | 0.325 | -0.66 | *p* < 0.001 |
| Friendliness | 0.16 | 0.387 | -0.70 | *p* < 0.001 | 0.64 | *p* < 0.001 | -0.03 | 0.882 |
| Anxiety | -0.24 | 0.208 | 0.32 | 0.089 | -0.58 | *p* < 0.001 | 0.14 | 0.472 |
| ***Rhesus macaque (N = 30: [25])*<sup>(2)</sup>** |  |  |  |  |  |  |  |  |
| Confidence | 0.76 | *p* < 0.001 | -0.34 | 0.069 | 0.38 | 0.038 | 0.11 | 0.561 |
| Openness | -0.10 | 0.603 | -0.19 | 0.311 | 0.00 | 0.999 | -0.05 | 0.792 |
| Dominance | 0.55 | 0.002 | 0.61 | *p* < 0.001 | -0.04 | 0.824 | -0.12 | 0.523 |
| Friendliness | 0.31 | 0.090 | -0.27 | 0.147 | 0.79 | *p* < 0.001 | -0.27 | 0.144 |
| Activity | 0.11 | 0.564 | 0.25 | 0.177 | 0.16 | 0.408 | -0.72 | *p* < 0.001 |
| Anxiety | -0.17 | 0.365 | 0.72 | *p* < 0.001 | -0.24 | 0.202 | -0.28 | 0.136 |
| ***Barbary macaque (N = 30: [22])*<sup>(2)</sup>** |  |  |  |  |  |  |  |  |
| Confidence | 0.84 | *p* < 0.001 | -0.25 | 0.175 | 0.46 | 0.010 | 0.01 | 0.958 |
| Openness | -0.23 | 0.224 | 0.31 | 0.097 | -0.28 | 0.128 | -0.09 | 0.636 |
| Friendliness | 0.26 | 0.170 | 0.23 | 0.223 | 0.55 | 0.001 | -0.48 | 0.007 |
| Irritability | 0.08 | 0.681 | 0.73 | *p* < 0.001 | -0.46 | 0.011 | -0.05 | 0.807 |
| ***Assamese macaque (N = 30: [22])*<sup>(2)</sup>** |  |  |  |  |  |  |  |  |
| Confidence | 0.84 | *p* < 0.001 | -0.20 | 0.296 | 0.26 | 0.163 | -0.01 | 0.956 |
| Activity | -0.04 | 0.846 | 0.51 | 0.004 | 0.23 | 0.218 | -0.68 | *p* < 0.001 |
| Openness | 0.19 | 0.308 | 0.49 | 0.007 | -0.17 | 0.383 | -0.25 | 0.184 |
| Friendliness | 0.20 | 0.300 | -0.34 | 0.070 | 0.74 | *p* < 0.001 | -0.14 | 0.450 |
| Opportunism | 0.57 | *p* < 0.001 | 0.57 | *p* < 0.001 | 0.04 | 0.824 | -0.04 | 0.828 |
| ***Tonkean macaque (N = 30: [22])*<sup>(2)</sup>** |  |  |  |  |  |  |  |  |
| Dominance | 0.83 | $p < 0.001$ | 0.16 | 0.392 | 0.18 | 0.331 | 0.01 | 0.938 |
| Openness | -0.06 | 0.757 | 0.55 | 0.002 | 0.10 | 0.614 | -0.58 | $p < 0.001$ |
| Friendliness | 0.01 | 0.952 | -0.53 | 0.003 | 0.62 | $p < 0.001$ | 0.04 | 0.836 |
| Sociability | -0.04 | 0.838 | -0.07 | 0.704 | 0.61 | $p < 0.001$ | -0.12 | 0.542 |
| ***Crested macaque (N = 30: [22])***<sup>(2)</sup> |  |  |  |  |  |  |  |  |
| Friendliness | 0.18 | 0.335 | -0.14 | 0.472 | 0.57 | 0.001 | -0.36 | 0.052 |
| Confidence | 0.80 | $p < 0.001$ | -0.23 | 0.228 | 0.40 | 0.027 | -0.12 | 0.511 |
| Aggressiveness | 0.36 | 0.053 | 0.83 | $p < 0.001$ | -0.09 | 0.644 | -0.32 | 0.082 |
| Excitability | 0.15 | 0.427 | -0.01 | 0.973 | -0.02 | 0.904 | -0.11 | 0.570 |
| ***Orangutan (N = 27: [51])***<sup>(2)</sup> |  |  |  |  |  |  |  |  |
| Extraversion | -0.06 | 0.763 | 0.06 | 0.757 | 0.39 | 0.042 | -0.58 | 0.001 |
| Dominance | 0.67 | $p < 0.001$ | 0.57 | 0.002 | -0.02 | 0.925 | -0.05 | 0.812 |
| Neuroticism | -0.61 | $p < 0.001$ | 0.64 | $p < 0.001$ | -0.35 | 0.075 | -0.32 | 0.102 |
| Agreeableness | 0.09 | 0.658 | -0.44 | 0.023 | 0.74 | $p < 0.001$ | -0.19 | 0.351 |
| Intellect | 0.39 | 0.044 | -0.12 | 0.541 | 0.01 | 0.972 | -0.11 | 0.573 |
| ***Chimpanzee (N = 26: [50])***<sup>(2)</sup> |  |  |  |  |  |  |  |  |
| Dominance | 0.82 | $p < 0.001$ | 0.15 | 0.477 | 0.26 | 0.196 | -0.06 | 0.757 |
| Extraversion | 0.24 | 0.236 | -0.17 | 0.412 | 0.51 | 0.008 | -0.53 | 0.006 |
| Conscientiousness | -0.21 | 0.302 | -0.80 | $p < 0.001$ | 0.32 | 0.114 | 0.14 | 0.510 |
| Agreeableness | 0.03 | 0.892 | -0.70 | $p < 0.001$ | 0.61 | $p < 0.001$ | 0.12 | 0.555 |
| Neuroticism | 0.02 | 0.910 | 0.59 | 0.002 | 0.11 | 0.606 | -0.34 | 0.089 |
| Openness | -0.22 | 0.273 | 0.03 | 0.877 | -0.32 | 0.109 | -0.43 | 0.030 |
| ***capuchin monkey (N = 30: [24])***<sup>(2)</sup> |  |  |  |  |  |  |  |  |
| Assertiveness | 0.77 | $p < 0.001$ | 0.35 | 0.059 | 0.15 | 0.434 | -0.07 | 0.720 |
| Openness | 0.36 | 0.051 | -0.11 | 0.574 | 0.31 | 0.101 | -0.59 | $p < 0.001$ |
| Neuroticism | -0.38 | 0.037 | 0.82 | $p < 0.001$ | -0.34 | 0.067 | -0.39 | 0.032 |
| Sociability | 0.25 | 0.175 | -0.27 | 0.145 | 0.70 | $p < 0.001$ | -0.14 | 0.446 |
| Attentiveness | 0.14 | 0.461 | -0.45 | 0.012 | 0.43 | 0.018 | -0.38 | 0.037 |

<sup>(1)</sup> Dom<sub>PR-2</sub> = Dominance<sub>PR-2</sub>; Agg<sub>PR-2</sub> = Aggressiveness<sub>PR-2</sub>; Soc<sub>PR-2</sub> = Sociability<sub>PR-2</sub>; Act<sub>PR-2</sub> = Activity<sub>PR-2</sub>
<sup>(2)</sup> *N* indicates the number of items that were common to the rating items used in this study.

### Personality structure of behavioural measures

*ICCs* suggested that 15 of the 26 behavioural measures were temporally stable (S8 Table); therefore, we conducted principal component analysis for 15 behavioural variables (Table 6). Thirteen of the 15 variables were transformed by the square root or logarithm to achieve normal distribution (S9 Table). Four components with eigenvalues exceeded 1.00. Examination of the scree plot suggested that there were four components, and parallel analysis indicated three components. To correspond with the rating results, we extracted four components and calculated the principal component scores based on varimax solution. We labelled the four behavioural components: Kin-biased-approaching_BO_, Grooming-diversity_BO_, Activity_BO_, and Aggressiveness_BO_ (Table 6). These four components accounted for 75 % of the total variance. The average absolute value of Pearson’s correlation coefficient between component scores and the subject’s attributes (Table 2) was *r* = 0.37 (SD = 0.29, range 0.07–0.68) for subject’s dominance rank, *r* = 0.21 (SD = 0.09, range 0.14–0.34) for age, and *r* = 0.31 (SD = 0.31, range 0.02–0.74) for the total number of kin.

**Table 6.** Factor loadings, eigenvalues, and cumulative contribution ratio of four components calculated by principal component analysis (varimax rotation) of behavioural data (N = 27).

| Behavioural measures | Components<sup>(1)</sup> |  |  |  |
| --- | --- | --- | --- | --- |
|  | Kba<sub>BO</sub> | Grd<sub>BO</sub> | Act<sub>BO</sub> | Agg<sub>BO</sub> |
| Rate approaching kin (within 3 m and arm-reach) | **0.93** | 0.23 | -0.08 | 0.03 |
| Kin-biased approaching | **0.87** | -0.06 | 0.25 | 0.01 |
| Prop time spent resting | **-0.68** | -0.23 | 0.28 | 0.26 |
| Prop time spent in quasi-proximity (3 m, arm-reach, and contact) | **0.42** | -0.06 | -0.30 | **-0.40** |
| Rate approaching within arm-reach (60 cm) | **0.60** | **0.68** | -0.24 | 0.18 |
| Kin-biased grooming | **0.43** | **-0.69** | -0.12 | 0.25 |
| Shannon-Wiener's diversity index | 0.13 | **0.95** | 0.10 | 0.02 |
| Number of grooming partners | 0.24 | **0.92** | -0.03 | 0.00 |
| Rate receiving submissive behaviour | 0.23 | -0.13 | **0.75** | 0.09 |
| Prop time spent moving | 0.01 | -0.03 | **0.74** | **0.49** |
| Prop time spent lying down | 0.19 | -0.14 | **-0.74** | -0.16 |
| Rate approaching (within 3 m) | **0.54** | 0.01 | **-0.70** | 0.00 |
| Rate threats | -0.15 | 0.35 | **0.57** | **-0.52** |
| Rate submissive behaviour | -0.08 | 0.11 | -0.12 | **-0.80** |
| Prop time spent on the tree | -0.35 | 0.22 | 0.27 | **0.63** |
| Eigenvalues | 3.42 | 3.02 | 2.88 | 1.91 |
| Cumulative contribution | 0.23 | 0.43 | 0.62 | 0.75 |

Salient loadings $\geq |.4|$ are shown in **bold**.
<sup>(1)</sup> Kba<sub>BO</sub> = Kin-biased-approaching<sub>BO</sub>, Grd<sub>BO</sub> = Grooming-diversity<sub>BO</sub>, Act<sub>BO</sub> = Activity<sub>BO</sub>, and Agg<sub>BO</sub> = Aggressiveness<sub>BO</sub>

### Comparison of personality structure between ratings and behavioural measures

Three of the four components in Wave 2 were related to the components extracted from behavioural data (Table 7). Moderate associations were found between Sociability_PR-2-1_ and Activity_BO_ (*r* = −0.58, *p* < 0.01) and between Activity_PR-2-1_ and Kin-biased-approaching_BO_ (*r* = − 0.57, *p* < 0.01). Weak relationships were found between Aggressiveness_PR-2-1_ and Grooming-diversity_BO_ (*r* = 0.44, *p* < 0.05), Sociability_PR-2-1_ and Kin-biased-approaching_BO_ (*r* = −0.48, *p* < 0.05), and Activity_PR-2-1_ and Grooming-diversity_BO_ (*r* = −0.39, *p* < 0.05). Dominance_PR-2-1_ and Aggressiveness_BO_ did not correlate with any of the components.

**Table 7.** Pearson correlation coefficients (*r*) and *p*-values between components of personality ratings and behavioural observation (N = 27).

| Components | *Wave 2 ratings<sup>(1)</sup>* |  |  |  |  |  |  |  |
| --- | --- | --- | --- | --- | --- | --- | --- | --- |
|  | Dom<sub>PR-2-1</sub> |  | Agg<sub>PR-2-1</sub> |  | Soc<sub>PR-2-1</sub> |  | Act<sub>PR-2-1</sub> |  |
| | $r$ | $p$-value | $r$ | $p$-value | $r$ | $p$-value | $r$ | $p$-value |
| ***Components of behavioural measure*** |  |  |  |  |  |  |  |  |
| Kin-biased-approaching<sub>BO</sub> | -0.12 | 0.540 | 0.07 | 0.725 | -0.48 | 0.011 | -0.57 | 0.002 |
| Grooming-diversity<sub>BO</sub> | -0.29 | 0.139 | 0.44 | 0.023 | 0.24 | 0.222 | -0.39 | 0.042 |
| Activity<sub>BO</sub> | 0.09 | 0.644 | 0.35 | 0.074 | -0.58 | 0.002 | -0.02 | 0.927 |
| Aggressiveness<sub>BO</sub> | 0.04 | 0.843 | 0.06 | 0.755 | 0.05 | 0.792 | 0.18 | 0.356 |

<sup>(1)</sup> Dom<sub>PR-2-1</sub> = Dominance<sub>PR-2-1</sub>; Agg<sub>PR-2-1</sub> = Aggressiveness<sub>PR-2-1</sub>; Soc<sub>PR-2-1</sub> = Sociability<sub>PR-2-1</sub>; Act<sub>PR-2-1</sub> = Activity<sub>PR-2-1</sub>

## Discussion

### Comparison of personality structure between Japanese macaques and the other non-human primates

Principal component analysis of Wave 2 ratings revealed four components: Dominance_PR-2_, Aggressiveness_PR-2_, Sociability_PR-2_, and Activity_PR-2_. Factors similar to Dominance_PR-2_ have been found in other non-human primate studies that used personality ratings (Table 5). In rhesus macaques, Assamese macaques, orangutans, and capuchin monkeys, Dominance_PR-2_ was strongly correlated with two factors. This may be because previous studies extracted more than four factors [25]. By specifying a larger number of factors in the analysis, what would originally be extracted as one factor might be separated into two factors which are defined more specifically.

Traits similar to Aggressiveness_PR-2_ in this study were found in other non-human primates (Table 5). Irritability of Barbary macaques contains the items: *sympathetic*, *protective*, and *helpful*; Neuroticism of capuchin monkeys includes the items: *Decisive* and *Impulsive*, and these items were not included in Aggressiveness_PR-2_ of Japanese macaques. Aggressiveness_PR-2_ may be more similar to Irritability of Barbary macaques and to Neuroticism of capuchin monkeys when adding the aspect of Big Five Agreeableness to Aggressiveness_PR-2_ of Japanese macaques. Aggressiveness_PR-2_ was also associated with Conscientiousness, Agreeableness, and Neuroticism in chimpanzees (Table 5). This result is consistent with that of Weiss et al. [25]. Chimpanzee Conscientiousness could be referred to as ‘individual differences in ability to control excitement,’ Agreeableness as ‘individual differences in trying to direct excitement to other individuals or not,’ Neuroticism as ‘individual differences in whether or not it is easily excitable due to its anxious tendency.’ This may be similar to the results of human studies in which Neuroticism and Agreeableness were associated with aggressive behaviour [52], and further research is needed to clarify what these results imply.

Sociability_PR-2_ in this study was similar to Friendliness and Sociability of other macaques and capuchin monkeys (Table 5). Sociability_PR-2_ was associated with Confidence, Friendliness, and Irritability of Barbary macaques and Friendliness and Confidence of crested macaques. Barbary and crested macaque Confidence were also highly correlated with Dominance_PR-2_ in this study. Barbary and crested macaques are egalitarian species compared to Japanese macaques [16]. Due to the obscurity of dominance rank relationships in egalitarian species, Sociability_PR-2_ in Japanese macaques may have been related to both Friendliness and Confidence.

Activity_PR-2_ in this study was similar to Activity or Openness in other non-human primates (Table 5). This was because Activity_PR-2_ contained items such as *Active* and *Lazy*. However, Activity_PR-2_ was not related to Openness in rhesus macaques from a previous study [25]. This may be because the ratings were effected by differences in age compositions of the two groups [53]. The average age of Wave 2 subjects in this study was 21.38 years (SD = 4.09 years, aged 13–31 years; S1 Table), whereas the average age of rhesus macaque subjects in the previous study [25] was 7.71 years (SD = 6.21 years). Factors such as macaque Activity and Openness found in previous studies may have different results, depending on the age.

Based on these comparisons of personality structure (Table 5), factors common to macaques and capuchin monkeys could be classified into the following four categories: (1) “dominance” factor which correlates with dominance rank, (2)"excitability” factor that can be described as Aggression or Neuroticism, (3) “sociability” factor that indicates how actively one individual is trying to interact with the others, and (4) degree of “activity” often exhibited as exploratory behaviour or Openness.

Some researchers have noted that biases in analysis could occur due to assessment methods; for example, “jingle-jangle fallacies” [54]. Cross-species comparisons could also include jingle-jangle fallacies or biases associated with attribution of factors [32]. The strong correlation between Aggressiveness_PR-2_ in this study and Friendliness in the other group of Japanese macaques (Table 5) indicated that these traits were the same but had different names (jangle fallacy). Stereotypical biases of dominance rank relationships may have affected the results regarding Dominance-related factors being found in any non-human primate during the interspecies comparison (Table 5). Simplified representations of observable behaviour [32] and the implicit semantic structure contained in ordinary language may have projected onto conclusions regarding personality structure [55].

### Temporal stability of personality structure

Temporal stability or instability in personality structure cannot reflect an intrinsic property but rather be a consequence of external environmental fluctuations [40,56]. The components of Dominance_PR-1_, Dominance_PR-2_, Kin-biased-approaching_BO_, and Activity_BO_ in this study were correlated with dominance rank or the number of kin (Table 2). These results suggest that the scales or measures constituting these components were temporally stable as they reflected the stable social structure of dominance rank or number of kin. Future studies will need to determine the situations in which temporal stability is observed and whether macaques have personality coherence. For example, rank order correlations could be found in measurements associated with stress-induced behaviour under certain conditions, such as agonistic interactions, and there might be individual differences in whether individuals are part of frequent agonistic interactions.

### Personality structure of behavioural measure

Our finding that Kin-biased-approaching_BO_ was positively associated with the number of kin (Table 2) was similar to that of a previous study on wild Chacma baboons in which the trait labelled Loner was correlated with the presence of kin [20]. High Kin-biased-approaching_BO_ individuals tended to be close to other individuals when they rested (Table 6). Kin-biased-approaching_BO_ and Activity_BO_ correlated with the number of kin or dominance rank (Table 2). High-ranking individuals stay around the provisioning site, and low-ranking individuals are around peripheral areas [57,58], and Kin-biased-approaching_BO_ and Activity_BO_ exhibited these in the subject’s behavioural tendencies. These results suggest that when the traits are assessed by observing behaviour which regularly occurs in wild populations, it may be easier to find traits that reflect behavioural characteristics associated with the social structure of macaques.

Kin-biased-approaching_BO_ has not been reported by previous studies. The reason for this was that other studies did not include behavioural measures about the macaques relationship with their kin. Because Japanese macaques have a stricter dominance rank hierarchy [16] and strong connections between kin [12–14] than other macaques such as crested macaques, Kin-biased-approaching_BO_ might have been easier to construct.

Previous studies have shown that macaques have close relationships with unrelated individuals, even if there are kin in the group, and grooming diversity is not associated with the number of kin or dominance rank [57,59]. Grooming-diversity_BO_ in this study was consistent with that in previous studies (Table 2). In addition, Grooming-diversity_BO_ appears to be similar to Sociability in crested macaques [19], which is composed of behavioural measures such as *diversity grooming partners*. Sociability was pointed out as a trait specific to crested macaques that is considered more tolerant than other macaques [19], but this study also found a component similar to Sociability, indicating that the trait of Grooming-diversity_BO_ might be a common trait of macaques.

Individuals with low Aggressiveness_BO_ are less likely to excite and more likely to be on the tree (Table 6), and this component was associated with dominance rank (Table 2) and submissive behaviour but not with receiving submissive behaviour (Table 6). These results may indicate that low-ranking individuals may be less likely to engage in agonistic interactions as they try to stay away from aggressive high ranks. If this is the case, it might be that non-human primates evaluate each other’s aggressiveness and adjust their behavioural patterns [20] so that they do not get involved in stressful situations such as agonistic interactions. Thus, the elucidation of how non-human primates perceive the personality of other members of the group will need to be explored further, as primate personality research progresses.

### Comparison of Japanese macaque’s personality structure between personality ratings and behavioural data

A comparison of Japanese macaques’ personality structure between personality ratings and behavioural data revealed that different personality structures were found in the two methods (Table 7). Activity_PR-2-1_ (from both Wave 1 and 2) was associated with Kin-biased-approaching_BO_, and Sociability_PR-2-1_ was associated with Activity_BO_, possibly because both traits reflected behaviour related to rank relationship of the Arashiyama group (see ‘Personality structure of behavioural measure’ section).

Personality structure of rating data did not reflect component structure found in the behavioural data (Table 7); that is, behavioural observations and personality ratings assessed different aspects of Japanese macaque behaviour [60]. We suggest at least three reasons regarding why different personality structures were found between the two methodologies:

1. Presence or absence of behavioural measures that cover a certain trait in ratings in the research design: Differences in factor structure are influenced by composition of the scale rather than reflecting potential specific properties [61]. When observations do not contain behavioural variables, such as novelty seeking, that can be measured through experiments, factor analysis of behavioural data in natural situations could negate the effect of a particular situation on behaviour and reveal a structure that is different from personality ratings. When this is the case, it could imply behavioural plasticity, as situational consistency of behaviour is a prerequisite for this assumption.
2. Problems with the personality rating methodology [33]: Previous studies have shown that some attribution bias occurs when explaining behaviour using existing information. For example, stereotypical biases such as age, sex, and rearing-related differences influence assessments [32]. People have simplified and made consistent representations of mental images for observable behaviours [32,39]. This is consistent with the results of strong temporal stability in Dominance_PR-1_ of this study (Table 4).
3. Personality coherence: That is, individuals may alter their behaviour based on the situation at any time [39]. For example, individuals may avoid stressful situations where conflicts can occur or they may stay away from other aggressive individuals[20]. Behavioural measures such as aggressive behaviours may not easily occur and may not be detected as temporally stable behaviours (S8 Table) when behavioural observations are conducted in wild populations.

It is the first-person perspective based on self-assessment and the second-person perspective based on interpersonal communication that creates temporal and cross-situational consistency of personality, and no consistency is detected from the third-person perspective data (for example, behavioural data) [37]. This idea of the second- and third-person perspectives may also be applicable to personality ratings and behavioural observation. This effect might be significant, especially in the relationship between zoo animals and their keepers, but it is not clear from our data. There is no research on such actors’ perspectives and animal personalities, and further research is needed to elucidate the same.

### Limitations of this study

This study had three methodological limitations. First, the short duration of observation time (mean per subject ± SD = 7.27 ± 0.61 h) may have affected the results. For example, previous studies have used self-grooming and scratching as behavioural measures of anxiety; however, in this study, these behavioural indicators had no temporal stability. There was also no temporal stability in aggressive behaviour in this study. It is possible that the number of observed cases was insufficient to assess temporal stability, because the behaviour was coded in a wide variety of situations under natural settings. Second, the sample size of this study was small for a study of the type that describes personality by identifying trait factors. This limited the ability to generalise results. Third, there is a possibility that Type I error may have occurred in the analysis. In this study, we used correlation coefficients to compare the personalities of Japanese macaques and other primates, and structure of personality was based on ratings and behavioural observations. Although *p*-values were reported for all correlation coefficients, the correlation coefficients in the results may have been reported to be higher by chance. In the future, clarifying the above methodological issues will help report more robust data.

## Acknowledgments

We are grateful for support provided by Mr. S. Asaba, all the staff at Iwatayama Monkey Park, and the raters who collected data. We would like to thank the anonymous referees for their constructive criticism of this manuscript. We thank Y. Gondou, M. Ueno, and N. Katsu for their insightful comments on the study.

## Supporting Information

**S1 Table**. Individual information.

**S2 Table**. Items used in Wave 1 personality ratings.

**S3 Table**. Items used in Wave 2 personality ratings.

**S4 Table**. Definitions of behavioural measures.

**S5 Table**. ICCs of items in Wave 1 ratings.

**S6 Table**. ICCs of items in Wave 2 ratings.

**S7 Table**. Personality structure in Wave 2 by three raters.

**S8 Table**. ICCs of 26 behavioural measures.

**S9 Table**. Behavioural data of principal component analysis.

**S10 Table**. Scores in Wave 1 and Wave 2 ratings.

**S11 Table**. Original behavioural data.

